# Characterization of opposing responses to phenol by *Bacillus subtilis* chemoreceptors

**DOI:** 10.1101/2021.08.31.458471

**Authors:** Girija A. Bodhankar, Payman Tohidifar, Zachary L. Foust, George W. Ordal, Christopher V. Rao

**Author notes:** To whom correspondence should be addressed: Christopher V. Rao, Department of Chemical and Biomolecular Engineering, University of Illinois at Urbana-Champaign, Urbana, Illinois, 61801, USA.

## Abstract

*Bacillus subtilis* employs ten chemoreceptors to move in response to chemicals in its environment. While the sensing mechanisms have been determined for many attractants, little is known about the sensing mechanisms for repellents. In this work, we investigated phenol chemotaxis in *B. subtilis*. Phenol is an attractant at low, micromolar concentrations, and a repellent at high, millimolar concentrations. McpA was found to be the principal chemoreceptor governing the repellent response to phenol and other related aromatic compounds. In addition, the chemoreceptors McpC and HemAT were found to the govern the attractant response to phenol and related compounds. Using receptor chimeras, McpA was found to sense phenol using its signaling domain rather than its sensing domain. These observations were substantiated *in vitro,* where direct binding of phenol to the signaling domain of McpA was observed using saturation-transfer difference nuclear magnetic resonance. These results further advance our understanding of *B. subtilis* chemotaxis and demonstrate that the signaling domain of *B. subtilis* chemoreceptors can directly sense chemoeffectors.

**IMPORTANCE:** Bacterial chemotaxis is commonly thought to employ a sensing mechanism involving the extracellular sensing domain of chemoreceptors. Some ligands, however, appear to be sensed by the signaling domain. Phenolic compounds, commonly found in soil and root exudates, provide environmental cues for soil microbes like *Bacillus subtilis*. We show that phenol is sensed both as an attractant and a repellent. While mechanism for sensing phenol as an attractant is still unknown, we found that phenol is sensed as a repellent by the signaling domain of the chemoreceptor McpA. This study furthers our understanding of the unconventional sensing mechanisms employed by the *B. subtilis* chemotaxis pathway.

## INTRODUCTION

Flagellated bacteria can swim up gradients of chemicals favorable to their growth, known as attractants, and down ones inhibitory to their growth, known as repellents (1). They sense these chemicals using chemoreceptors. *Bacillus subtilis*, a Gram-positive soil bacterium, possesses ten chemoreceptors. Eight are transmembrane and two are soluble (2). The transmembrane chemoreceptors possess an extracellular sensing domain and an intracellular signaling domain. The extracellular sensing domain is coupled to the intracellular signaling domain by two transmembrane helices, known as TM1 and TM2, and an intracellular HAMP domain (3). The soluble chemoreceptors also have signaling and sensing domains but lack the transmembrane helices and HAMP domain.

The *B. subtilis* chemoreceptors form stable complexes with the histidine kinase CheA and coupling proteins CheV and CheW (4, 5). These complexes form large, hexagonally structured clusters that preferentially localize at the poles of the cell (6, 7). These clusters are able to amplify the signaling response through allosteric interactions with neighboring chemoreceptors (8). The binding of attractants to the *B. subtilis* chemoreceptors increases the rate of CheA autophosphorylation (5). The phosphoryl group is then transferred to CheY, a soluble response regulator. Phosphorylated CheY can then bind the flagellar motors and induces them to spin counter clockwise, which causes *B. subtilis* to swim in a straight direction (5, 9). Presumably, the binding of repellents to *B. subtilis* chemoreceptors decreases the rate of CheA autophosphorylation. This, in turn, decreases the concentration of phosphorylated CheY, which reduces the likelihood that the motors spin counter clockwise. When not bound with phosphorylated CheY, the motors spin clockwise, which causes *B. subtilis* to tumble about. Chemical gradients are sensed through a temporal mechanism involving sensory adaptation. *B. subtilis* uses three complementary mechanisms for sensory adaptation involving receptor methylation, allosteric regulation by a soluble protein known as CheD, and phosphorylation of the CheV adaptor protein (4, 10, 11).

Multiple studies have investigated the mechanisms for sensing attractants by chemoreceptors in *B. subtilis* and other diverse species of bacteria (see (12) for a comprehensive review). However, far less is known about the sensing mechanisms involving repellents. In this study, we investigated phenol chemotaxis in *B. subtilis*. Phenol is an attractant at low, micromolar concentrations, and a repellent at high, millimolar concentrations. By analyzing different mutants, McpA was found to be the principal chemoreceptor governing the repellent response to phenol and other related aromatic compounds. In addition, the chemoreceptors McpC and HemAT were found to the govern the attractant response though sensing mechanisms remain unknown. Using receptor chimeras, McpA was found to sense phenol using its signaling domain rather than its sensing domain. These observations were substantiated *in vitro,* where direct binding of phenol to the signaling domain of McpA was observed using saturation-transfer difference nuclear magnetic resonance.

## RESULTS

### *B. subtilis* senses phenol as an attractant at low concentrations and a repellent at high concentration

The chemotactic response to phenol and related aromatic compounds was measured using the capillary assay (1). In this assay, glass capillaries containing test compounds are introduced in wells filled with bacteria in chemotaxis buffer. The diffusion of these test compounds out of the capillary generates a chemical gradient. Bacteria that enter the capillary after 1 h of incubation are counted to quantify the chemotaxis response. The base response in the absence of a chemical gradient is determined using a control capillary containing buffer only. A colony count greater than the base count indicates an attractant response while a lower colony count indicates a repellent response. Phenol was chosen as the model compound to elucidate the chemotactic response to aromatic compounds. Phenolic compounds are abundant in nature, and chemotaxis to phenol has been studied in other bacteria (13).

The capillary assay response to increasing concentrations of phenol is shown in **Fig. 1A**. The number of bacteria accumulating in the capillaries was greater than the base count for capillaries containing phenol at concentrations less than 100 µM. A drop in the response was observed at higher concentrations. The decrease in response at higher concentrations could be due to a repellent response to phenol, loss of motility, or decreased viability of cells. Past studies have shown that the minimal inhibitory concentration of phenol for *B. subtilis* is 16 mM (14), suggesting a repellent response instead of decreased viability. To assess repellent responses, we used the “chemical-in-pond” modification of the capillary assay (15). In this modification, capillaries are filled with chemotaxis buffer and inserted in bacterial wells, called ponds, containing the repellent. The chemotaxis response is quantified as the number of bacteria entering the capillary. A potent repellent response to phenol was observed at a concentration of 1 mM in the pond (**Fig. 1A** **inset**). These results suggest that phenol is sensed as an attractant at low concentrations and a repellent at high concentrations.

**Figure 1.**
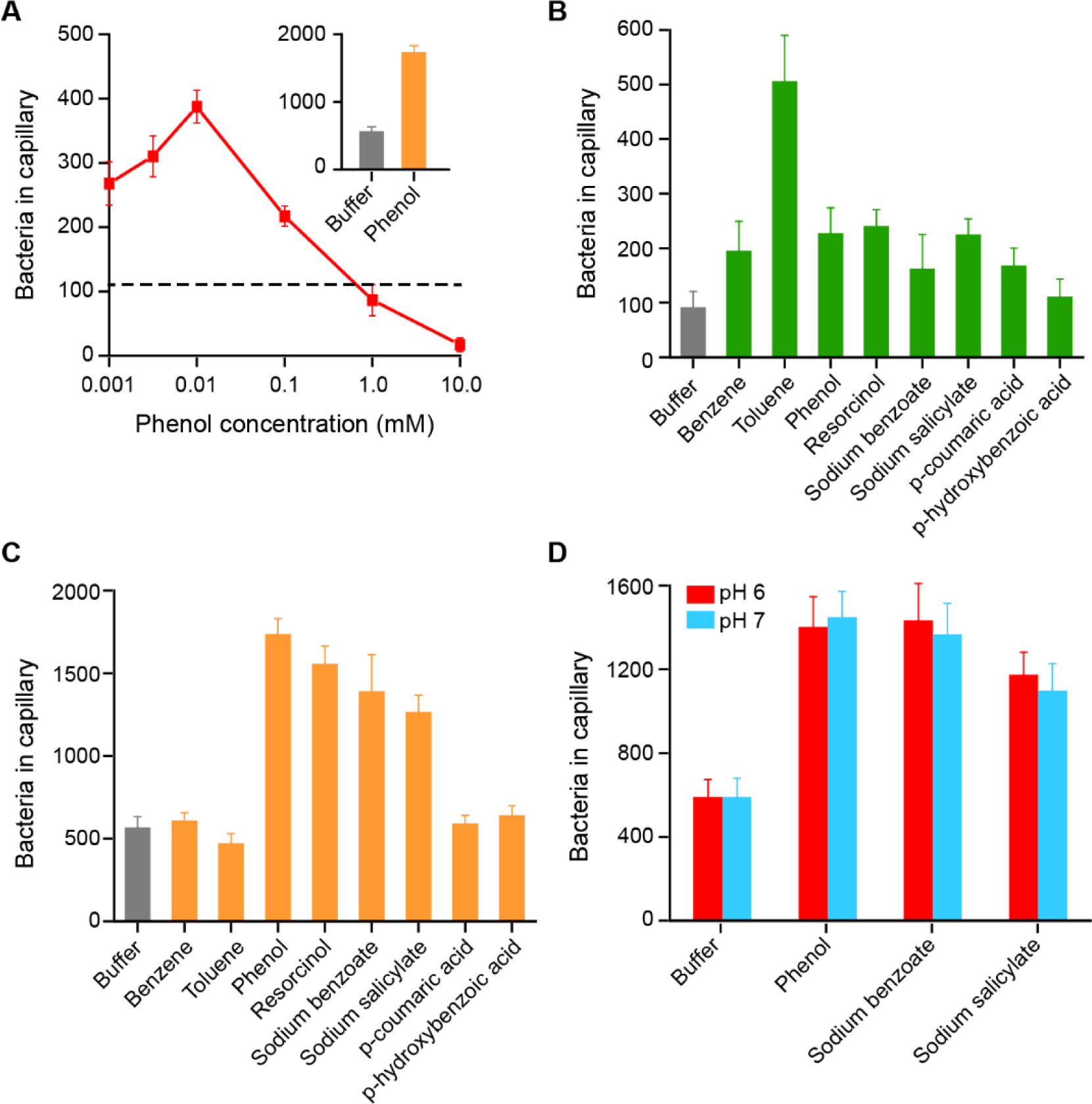
Attractant and repellent chemotaxis responses of wild-type *B. subtilis* towards phenol and other aromatic compounds. (A) Dose-dependent response of the wild-type strain to increasing concentrations of phenol measured using the attractant capillary assay. The dashed line indicates the base response to buffer control; inset shows the repellent response to 1 mM phenol measured in the repellent capillary assay. (B) Attractant chemotaxis response of the wild-type strain to 1 µM aromatic compounds in the attractant capillary assay. (C) Repellent chemotaxis response of the wild-type strain to 1 mM aromatic compounds in the repellent capillary assay. (D) Chemotaxis response of the wild-type strain to weak acids at neutral and pH 6.0 in the repellent capillary assay. Error bars indicate standard deviations obtained from three biological replicates performed at separate days.

To understand the ligand range and preference better, we tested the chemotactic responses to a number of aromatic compounds that differ in structures using the attractant and repellent capillary assays. Benzene and toluene were selected due to their simple structures. Physiologically relevant aromatics such as salicylate, benzoate, resorcinol, and p-hydroxybenzoate, which are found in plant root exudates, were also tested (16). Attractant assays were performed at a concentration of 1 µM in the capillary (**Fig. 1B**). Low concentrations were tested to avoid repellent effects (**Fig. 1B**). At 1 µM concentration, toluene had high attractant response. Other tested compounds except p-hydroxybenzoic acid had slightly higher responses than buffer. Repellent response was tested at 1 mM concentration in the pond (**Fig. 1C**). At this concentration, compounds that showed high repellent response were phenol, resorcinol, sodium benzoate, and sodium salicylate. A lower response was observed for benzene, toluene, p-coumaric acid, and p-hydroxybenzoic acid.

Benzoate and salicylate are known to induce weak repellent responses in *Escherichia coli* and *Salmonella enterica* at pH 5.5 (17, 18). When the external pH is lower than the internal pH, weak acids traverse the membrane to release protons and lower the cytoplasmic pH. To test whether the repellent response is due to a change in intracellular pH, capillary assay experiments were also carried out at a lower pH. The motility of *B. subtilis* is reduced at pH 5.5, thus the experiments were performed at an external pH of 6 (19). The repellent responses were unaffected due to the external pH change (**Fig. 1D**), suggesting that the response is not due to pH changes.

### McpA is the major chemoreceptor for sensing repellents

Many repellents are membrane active compounds, and the possibility that the repellent response is due to direct action on the membrane was dismissed in a previous study (20). The dose-dependent response of phenol in capillary assays also hinted at a receptor-specific response. *B. subtilis* has ten chemoreceptors (2). To identify the chemoreceptors involved in repellent sensing, we tested *B. subtilis* mutants expressing a single chemoreceptor with the other nine deleted (19) (**Fig. 2A**). Only the strain expressing McpA as its sole chemoreceptor exhibited a repellent response to phenol. To confirm this result, an *mcpA* knockout (Δ*mcpA*) mutant was tested. The Δ*mcpA* mutant’s repellent response to phenol was almost completely eliminated, confirming that McpA is the main repellent chemoreceptor (**Fig. 2A** **inset)**. Other chemoreceptors may sense phenol as a repellent but, at native expression levels, their contribution is minor and McpA dominates the response. Strains expressing McpA alone were also able to mediate a repellent response to other phenolic compounds tested in this study (**Fig. 2B**).

**Figure 2.**
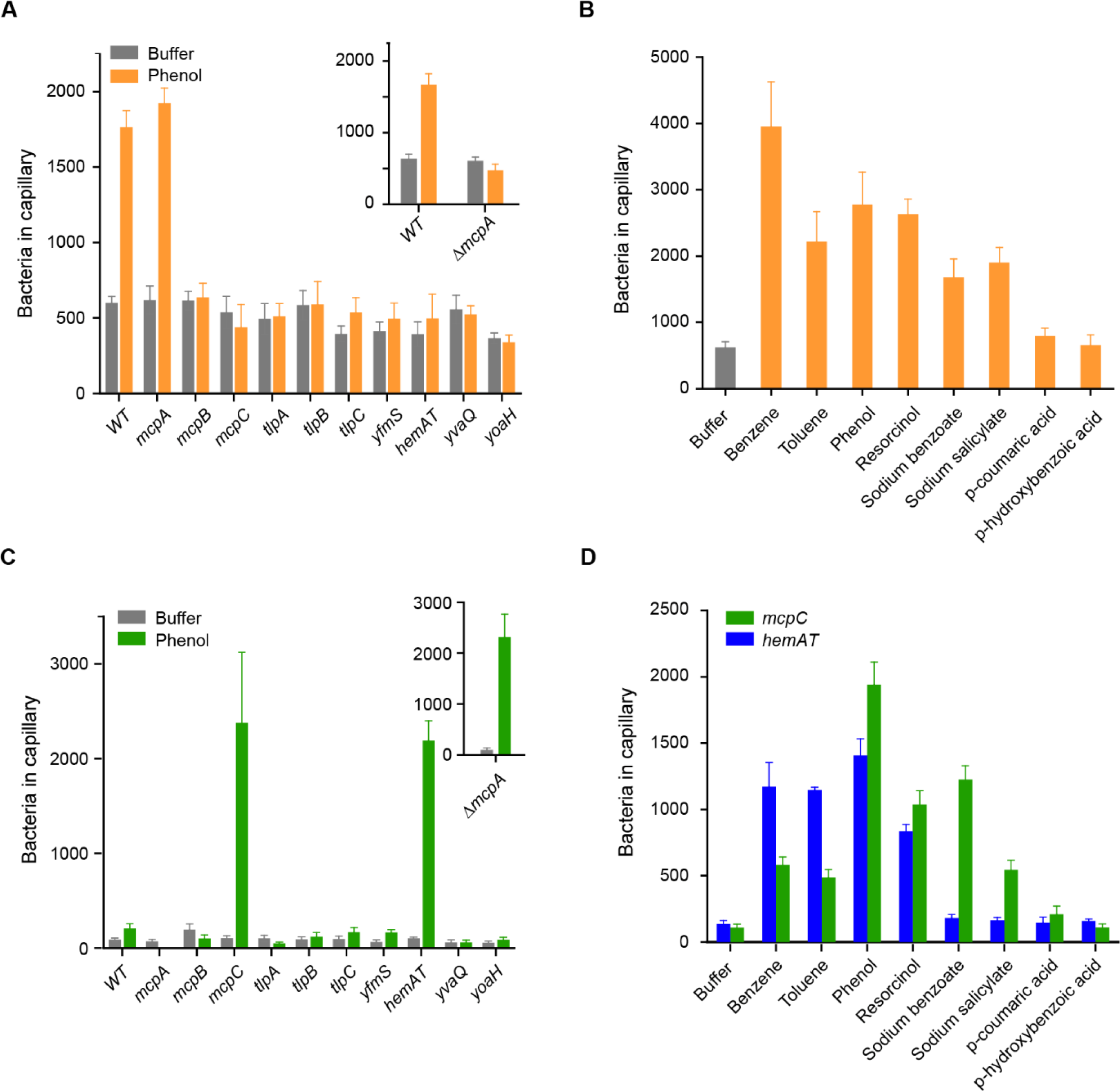
*B. subtilis* chemoreceptors involved in sensing phenol and other aromatic compounds as a repellent or an attractant. (A) Repellent chemotaxis responses of strains expressing single chemoreceptors to phenol measured in the repellent capillary assay; inset shows repellent chemotaxis response of a mutant lacking McpA chemoreceptor (Δ*mcpA*) to phenol. (B) Repellent chemotaxis responses of strains expressing McpA as their sole chemoreceptor to different aromatic compounds measured in the repellent capillary assay. (C) Attractant chemotaxis responses of strains expressing single chemoreceptors to phenol measured in the attractant capillary assay; inset shows attractant chemotaxis response of a mutant lacking McpA chemoreceptor (Δ*mcpA*) to phenol. (D) Attractant chemotaxis responses of strains expressing McpC or HemAT as their sole chemoreceptors to different aromatic compounds. Concentrations of chemoeffectors were set to 316 µM in the ponds and 0.1 M in the capillaries for repellent and attractant capillary assays, respectively. Error bars indicate standard deviations obtained from three biological replicates performed at separate days.

### Functional redundancy of attractant chemoreceptors

*B. subtilis* is attracted to phenol at low concentrations. In order to identify the chemoreceptors that sense phenol as an attractant, we tested single-receptor mutants using the attractant capillary assays (**Fig. 2C**). Mutant strains with McpC or HemAT as their sole chemoreceptors exhibited attractant responses to phenol. These responses were comparable to the control strain (Δ*mcpA)* deleted for the repellent response (**Fig. 2C** **inset).** It is notable that the attractant responses were not additive. Since the response to phenol at micromolar concentrations was low, higher concentrations (100 mM) of phenol were tested to compensate for the lack of signal amplification associated with allosteric interactions between chemoreceptors (21, 22). The strain with McpC as its sole chemoreceptor was able to sense other aromatic compounds as well, except p-coumaric acid and p-hydroxy benzoate (**Fig. 2D**). The strain with HemAT as its sole chemoreceptor sensed benzene, toluene, phenol, and resorcinol as attractants (**Fig. 2D**). McpC was previously found to sense amino acids and HemAT to sense oxygen and alcohols (23-25). Functional redundancy of chemoreceptors for sensing aromatic compounds is also observed in other bacteria (26).

### Cytoplasmic sensing of phenol

Transmembrane chemoreceptors have a modular structure consisting of an extracellular sensing domain and a conserved, intracellular signaling domain. They also have a cytoplasmic HAMP domain for relaying signals between the two domains (3). It was shown that phenol sensing by *E. coli* chemoreceptors is mediated by the transmembrane helices and HAMP domain (27). In order to determine the regions of McpA involved in sensing phenol as a repellent in *B. subtilis*, we used chimeric receptors generated by swapping domains between McpA and McpB (25, 28). McpB senses asparagine as an attractant using its extracellular sensing domain and is not involved in phenol sensing (29). This makes it a good control in order to confirm the functionality of the chimeras as well as to identify the regions of McpA involved in phenol sensing (**Fig. S1A**). All chimeras were expressed under the *mcpA* promoter in a strain deleted for the native chemoreceptors.

Strains expressing a McpA-McpB chimera with the sensing domain from McpA replaced with the one from McpB (*mcpA_44_B_267_A*) showed a repellent response to phenol and an attractant response to asparagine (**Fig. 3** and **Fig. S1A**). This suggests that the extracellular sensing domain of McpA receptor does not sense phenol. Likewise, strains expressing a McpA-McpB chimera with the McpB signaling domain replaced with the one from McpA (*mcpB_359_A*) showed a repellent response to phenol, indicating that the transmembrane helices and HAMP domain of McpA are not required for sensing phenol. The repellent response to phenol was eliminated in a chimera where the McpA signaling domain was swapped with the McpB signaling domain (*mcpA_358_B*). This mutant strain, however, responded to acidic pH mediated by McpA extracellular sensing domain (19), proving functionality of the chimera (**Fig. S1B**). These observations suggest that the repellent response to phenol is mediated by the intracellular signaling domain of McpA.

**Figure 3.**
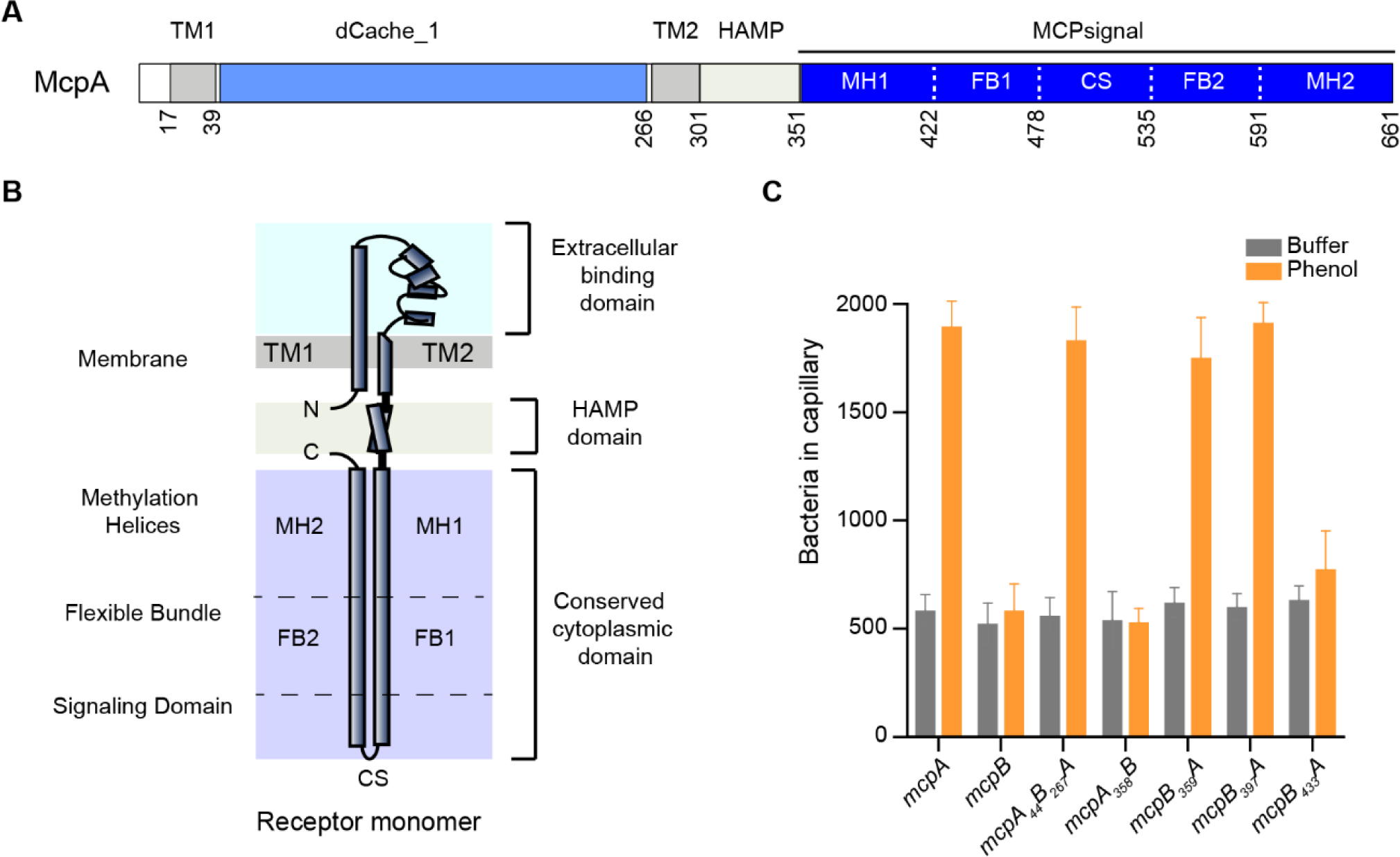
The signaling domain of McpA is involved in sensing phenol as a repellent. (A) Domain structure of McpA. The chemoreceptor consists of an extracellular sensing domain (dCACHE_1), two transmembrane helices (TM1 and TM2), followed by an intracellular HAMP domain, and an intracellular signaling domain (MCPsignal). The signaling domain is comprised of three subdomains known as the methylation helices (MH), flexible bundle (FB), and conserved signaling tip (CS). (B) Cartoon structure of McpA chemoreceptor monomer. (C) Repellent chemotaxis responses of strains expressing different chimeras between McpA and McpB as their sole chemoreceptor to 316 µM phenol measured in the repellent capillary assays. Error bars indicate standard deviations obtained from three biological replicates performed at separate days.

The signaling domains of chemoreceptors consist of coiled-coil helices with high sequence conservation across bacteria and archaea. They have three structurally distinct subdomains: the methylation helices, flexible bundle, and signaling tip (3). We tested receptor chimeras with fusions near the junction of the methylation helices and flexible bundle (*mcpB_397_mcpA* and *mcpB_433_*A). Cells expressing *mcpB_397_A* showed a repellent response to phenol while cells expressing *mcpB_433_A* failed to respond to phenol but responded to asparagine normally (**Fig. 3** and **Fig. S1A**). These results suggest that phenol is sensed by McpA using the region spanning residues 397 to 433, which corresponds to bottom of the methylation helix and top of flexible bundle on the N-terminus of McpA signaling of domain (see **Fig. 3**). Interestingly, the corresponding region on McpB is also involved in sensing short-chain aliphatic alcohols (C1-C5), namely ethanol (25).

Docking experiments confined to phenol-sensing region on McpA were carried out to gain some insights into putative residues involved in phenol binding. Computational predictions showed that phenol conformations with the lowest docking energy scores cluster within the dimer (ranging from Glu^397^to Ile^410^ on the N-helix and Glu^600^ to Ser^613^ on the C-helix). Interestingly, other repellent ligands that are sensed by McpA also clustered in the same region except for salicylate (**Fig. S2A** and **Fig. S2B**). Phenylalanine, an aromatic amino acid sensed by McpC, was used as a negative control for the docking experiments. Phenylalanine docked in a different region (residues Leu^417^ to Ser^430^ on the N-helix and Asp^579^ to Gln^593^ on the C-helix) (**Fig. S2C**). To assess whether these residues are important for phenol binding, we first aligned the amino-acid sequences spanning residues 397 to 410 on the N-helix and the neighboring residues 600 to 613 on the C-helix of McpA, McpB, TlpA, and TlpB (**Fig. S2D**). Note that these chemoreceptors are highly conserved at their signaling domain, and hence, were used to guide amino acid substitution experiments. Unfortunately, single substitution of non-conserved residues in McpA for the corresponding amino acids in McpB (N402K, A406S, S425A, A424T, M421V, S431A, S606A) did not show any change in response (data not shown).

The attractant chemoreceptors for phenol, McpC and HemAT, are phylogenetically distant from the other receptors in *B. subtilis* and, hence, not suitable for chimeric analysis in this study (**Fig. S3**). Indeed, chimeric receptors with the cytoplasmic domains swapped between McpA and McpC failed to respond to proline, which is sensed by the sensing domain of McpC (23). HemAT also proved to be unamenable to chimeric analysis using either McpA or YfmS (the only other soluble *B. subtilis* chemoreceptor). However, we were able to identify an aromatic attractant chemoeffector for McpB. Cells expressing McpB as their sole chemoreceptor were able to sense benzene as an attractant. We were further able to identify the region involved in sensing benzene as an attractant using chimeric receptors between McpA (repellent receptor for benzene) and McpB (attractant receptor for benzene). Strains expressing the chimera *mcpB_397_A*, involving the sensing domain of McpB and the signaling domain of McpA, showed a repellent response to benzene. However, strains expressing the chimera *mcpB_433_*A, which contains the previously mentioned 36 amino-acid region (residues 397 to 433) of the McpB signaling domain, showed an attractant response to benzene (**Fig. S4**), implying that the same region on McpB is also involved in sensing benzene as an attractant by McpB.

### *In-vitro* characterization of phenol-chemoreceptor interactions using STD-NMR

To determine whether the chemoreceptors directly bind phenol, we used ^1^H-STD-NMR (Proton Saturation-Transfer Difference Nuclear Magnetic Resonance) (30). STD-NMR is a small molecule-based NMR method for identifying ligands binding with medium to weak affinities (100 µM < K_d_ < 10 mM). This technique exploits the transfer of magnetization from a selectively irradiated protein to a ligand in its proximity (31). Ligand protons closest to the protein will receive magnetization more efficiently (32). As the protein-ligand complex is in equilibrium, the magnetization is transferred to the bulk solution upon ligand dissociation. This is observed as a decrease in the ligand peak in the NMR spectrum (on-resonance spectrum) over that of the reference spectrum with no selective saturation (off-resonance spectrum). The difference between the on- and off-resonance spectra will only have peak intensities corresponding to the ligand protons that bound to the protein.

**Fig. 4A** shows the ^1^H- and STD-NMR spectra for the signaling (McpAc) and the sensing (McpAs) domains of McpA incubated with 2 mM phenol. Phenol protons have resonance peaks at 7.2 ppm (meta), 6.9 ppm (para), and 6.8 ppm (ortho). Strong STD-NMR signals for all three proton groups were observed with the McpA signaling domain in the presence of phenol while STD-NMR signals for the McpA sensing domain incubated with the same concentration of phenol showed negligible peaks, corroborating chimeric receptor experiments. Additionally, no phenol peaks were observed for control experiment in the absence of proteins. To gain insight about the direction of phenol binding to the McpA signaling domain, we analyzed the STD amplification factor (STD-AF) for phenol protons. Briefly, the STD-AF for a proton is calculated as the ratio of integrated peak signal in the STD spectrum over the reference spectrum multiplied by ligand excess. A difference of at least 10% between ligand epitopes is recommended for classification as a preferred binding orientation (**Table 1**) (33). These analyses show that para proton has slightly higher STD effect compared to other protons (15% over ortho protons and 18.9% over meta protons), suggesting that the para proton is closer to the chemoreceptor. Competitive binding experiments have been used to distinguish between specific and non-specific binding in conjunction with STD-NMR (34, 35). However, these experiments require a reference ligand for which the binding site on protein and binding affinity between the two are known. Since such ligand was unknown for these experiments, we instead conducted the STD-NMR experiments for the McpB signaling domain in the presence of excess phenol to provide an additional negative control. The rational for this experimental design was that the McpB signaling domain has a coiled-coil structure with 74% sequence identity to the McpA signaling domain. As expected, negligible phenol peaks (STD-AF < 4%) were observed in the STD spectrum **(Fig. S5A)**. Finally, we conducted a control STD-NMR experiment with phenylalanine. Phenylalanine is an aromatic amino acid sensed by the McpC sensing domain with weak binding affinity (23). No signals corresponding to the ligand were observed in the STD spectrum of McpA signaling domain incubated with 2 mM phenylalanine (**Fig. S5B**). Chimeric receptor experiments suggested that the McpB signaling domain is involved in sensing benzene. STD spectrum of McpB signaling domain in the presence of 2 mM benzene showed a peak associated with benzene moiety with STD-AF of 16.4%, implying direct interaction of benzene with the McpB signaling domain (**Fig. S5C**).

**Figure 4.**
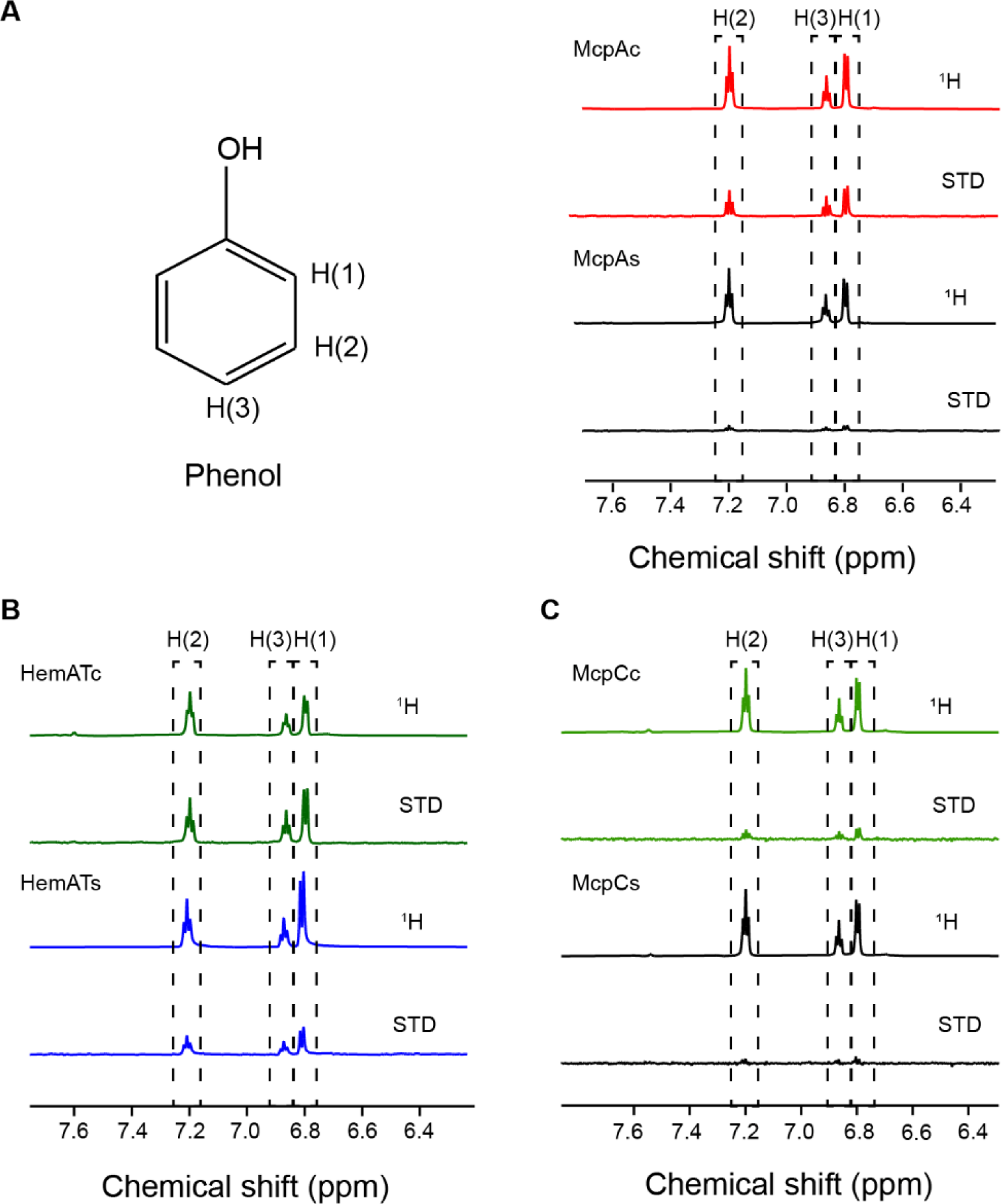
*In-vitro* characterization of binding between phenol and *B. subtilis* chemoreceptors. (A) ^1^H and STD-NMR spectra obtained from incubation of 20 µM McpA signaling (top) and sensing (bottom) domains with 2 mM phenol. (B) ^1^H and STD-NMR spectra obtained from incubation of 20 µM HemAT signaling (top) and sensing (bottom) domains with 2 mM phenol. (C) ^1^H and STD-NMR spectra obtained from incubation of 20 µM McpC signaling (top) and sensing (bottom) domains with 2 mM phenol. ^1^H peaks for ortho, meta, and para moieties of phenol are shown within dashed boxes.

**TABLE 1.** STD Amplification factors for phenol-protein interactions

| **Protein** | **H(2) Meta proton** | **H(3) Para proton** | **H(1) Ortho proton** |
| --- | --- | --- | --- |
| McpA<sub>C</sub> | 11.09 | 13.69 | 11.61 |
| McpA<sub>S</sub> | 3.52 | 5.54 | 3.72 |
| McpC<sub>C</sub> | 3.11 | 4.15 | 2.89 |
| McpC<sub>S</sub> | 0.82 | 1.33 | 1.28 |
| HemAT<sub>C</sub> | 23.85 | 26.90 | 24.29 |
| HemAT<sub>S</sub> | 17.41 | 19.50 | 15.87 |
| McpB<sub>C</sub> | 3.64 | 4.02 | 3.22 |

In the case of HemAT, phenol peaks were observed in the STD spectra for both the sensing and signaling domains (**Fig. 4B**). The para proton had the highest STD-AF values; however, the difference between STD-AF values within different moieties was too close to the 10% threshold described above. Therefore, it is difficult to classify this as a preferred binding direction. As negative controls, no binding was observed between phenylalanine and either HemAT domain. HemAT is a soluble chemoreceptor that contains a heme group for sensing molecular oxygen (36, 37). The sensing domain dimer interface consists of a four helical bundle at its core with other helices packed around it. HemAT sensing domain helices are hypothesized to be involved in sensing ethanol (25), and phenol likely follows a similar mechanism. The signaling domain of HemAT has a coiled-coil conserved structure similar to that of McpA. Surprisingly, the STD spectra for phenol and the McpC sensing and signaling domains were negligible (**Fig. 4C**). This suggests that phenol is probably sensed by the transmembrane helices or the HAMP domain of McpC.

We next used STD-NMR to determine the affinity of phenol for the signaling domain of McpA. STD-NMR titration experiments at different ligand concentrations can be used to determine the binding affinity (K_d_) of a protein-ligand complex. However, STD signals also rely on the kinetics of protein-ligand rebinding. The method of initial growth rates, where STD factors are calculated at the limit of zero saturation time (no protein-ligand rebinding), has been successfully used in an earlier report for determining K_d_ values (38). Briefly, STD-AF values were first calculated at different saturation times to get the initial-slope for each concentration (STD-AF0) (**Fig. 5A**). Initial slope values were then plotted against ligand concentrations to determine the K_d_ using the Langmuir isotherm curve. The binding affinity to phenol was calculated as 4.8 mM with McpA’s signaling domain (**Fig. 5B**). It is important to note that the binding curve for phenol and McpA signaling domain pair followed the hyperbolic curve model with a decreasing slope at higher phenol concentrations indicative of a specific binding event. However, at higher phenol concentrations (>4 mM), non-specific binding dominates, and the curve increased linearly instead of reaching a plateau. As a consequence, the K_d_ is overestimated in this study. The overestimation of binding K_d_ values has also been reported in another STD-NMR study and orthogonal techniques are recommended for obtaining accurate values (33, 38). As such, we conducted isothermal titration calorimetry experiments to assess the binding between McpA signaling domain and phenol. Unfortunately, these experiments were not conclusive because the affinity is weak (data not shown). In the case of HemAT, the phenol binding mechanism appears to be rather complex as phenol directly interacts with both sensing and signaling domains of HemAT.

**Figure 5.**
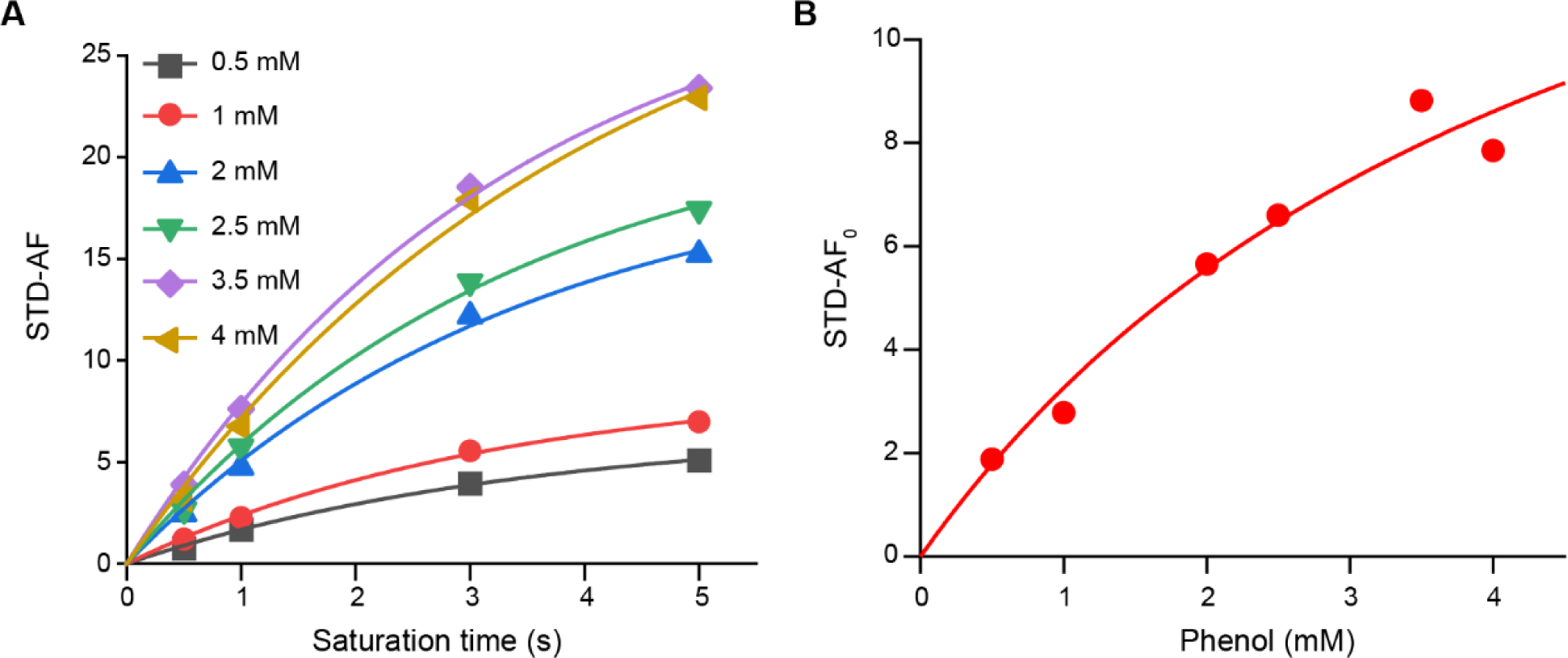
Estimation of dissociation constant (K_d_) of binding between the McpA signaling domain and phenol using STD-NMR. (A) Time evolution of STD amplification factor (STD-AF) values obtained from incubation of 20 µM McpA signaling domain with different concentrations of phenol (0.5 mM – 4 mM). The initial slope (STD-AF_0_) values were calculated from the fitted curves to estimate the binding dissociation constant. (B) STD-AF_0_ values from the previous step were fitted against different phenol concentrations to estimate the binding dissociation constant using the Langmuir isotherm model (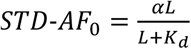; K_d_ = 4.8 mM, α = 18.9, R^2^=0.98).

## DISCUSSION

Previous studies reported several membrane-active agents are repellents for *B. subtilis* (39). Initially, it was unknown whether repellents are sensed by receptors or the membrane through changes in fluidity. Later, it was shown that repellents are sensed by chemoreceptors; however, their identities were unknown (40, 41). In this work, we found that the repellent receptor for phenol is McpA, a transmembrane chemoreceptor that also senses external acidic pH (19). McpA was found to be a broad-range chemoreceptor and capable of sensing multiple other aromatic compounds as repellents. Our data indicate that phenol is sensed by the cytoplasmic signaling domain of McpA. This contrasts the more common mechanism where chemoeffectors bind the extracellular sensing domain to induce signaling. That said, multiple examples of sensing by the signaling domain have been documented in the literature. For example, the region below the HAMP domain was also found to be important for sensing toluene and o-xylene by the Tar chemoreceptor in *E. coli* (42). Phenol sensing in *E. coli* also follows an unconventional route where phenol appears to bind the transmembrane helices and HAMP domain (27). In addition, the PctA chemoreceptor from *Pseudomonas aeruginosa* senses chlorinated compounds as repellents, but no binding was observed with the extracellular sensing domain of the chemoreceptor (43). Lastly, ethanol is sensed by the signaling domain of McpB in *B. subtilis* (25).

Chimeric receptor analysis revealed the importance of amino-acid residues near the junction of N-terminal methylation helices and flexible bundle in McpA. In case of attractant sensing, benzene also appears to be sensed by the same region in McpB. Swapping the 36 amino-acid regions between McpA and McpB chemoreceptors converted the repellent response of McpA to an attractant response, and vice versa for McpB. This region was also important for sensing ethanol by the signaling domain of McpB (25). Molecular dynamics simulations and mutation studies correlated changes in coiled-coil packing of the signaling domain with ability of McpB to sense ethanol. While the exact mechanism for signal propagation in chemoreceptors is still being resolved, kinase activity is thought to be modulated by changes in chemoreceptor dynamics that propagate the signal through HAMP and signaling domains (44-46). Recent studies have characterized chemoreceptor flexibility and dynamics in different signaling states. Electron paramagnetic resonance (EPR) studies have shown that the N-terminal methylation helices have different dynamics as compared to the C-terminal tail of the chemoreceptor in both methylated and demethylated states (47-49). Mobility of the N-terminal methylation helices was proposed to be a key signaling element in hydrogen exchange and solid-state NMR studies (50, 51). Cryoelectron tomography and molecular dynamics simulation studies of the *E. coli* Tsr chemoreceptor showed the importance of the glycine hinge (present in the flexible bundle region) in controlling chemoreceptor compactness and flexibility to allow for different signaling states (52, 53). It is possible that ligands like phenol and ethanol interact with signaling domain helices and cause changes in receptor packing or mobility, thus mimicking traditional signals that would have been propagated from the sensing domain via HAMP region. However, the exact mechanism of phenol sensing by McpA chemoreceptor remains unknown.

The attractant response to phenol is mediated by McpC and HemAT. McpC is a transmembrane chemoreceptor involved in sensing amino acids and sugars (23, 54). It is unclear how phenol induces signaling in McpC as no interaction between phenol and the sensing or signaling domains was observed by STD-NMR. These experiments fail to detect tight binding events (affinities in nanomolar-micromolar range) (38). Because the exchange of ligand is slow, the transfer to bulk solution is low and weak signals are observed. Another possibility is that phenol interacts with the transmembrane helices or HAMP domain of McpC in a manner similar to Tar and Tsr in *E. coli* (27); however, the possibility of an indirect sensing mechanism cannot be ruled out. The McpC sensing domain senses many amino acids indirectly though membrane associated proteins (23) while the signaling domain is involved in sugar taxis through interactions with the phosphotransferase system (PTS) (54). Thus, phenol could induce signaling through an indirect interaction. HemAT, on the other hand, is a cytoplasmic heme containing receptor known to sense molecular oxygen (24). Both the sensing and signaling domains of HemAT receptor bound phenol in the STD-NMR experiments. The sensing domain of HemAT is also involved in recognizing ethanol although the heme group is not involved (25). How phenol induces different output responses for different receptors is still an open-ended question.

Phenol is a complex chemoeffector and induces both attractant and repellent responses in *E. coli* and *S. enterica* (13, 55). The overall response is dependent on the relative abundance of different chemoreceptors (56). HemAT and McpA are the two most abundant receptors in *B. subtilis* (19,000 ± 3,900/cell and 15,900 ± 3,000/cell, respectively) while McpC chemoreceptor concentrations are low (2,800 ± 640/cell) (2). Unlike *E. coli* and *S. enterica,* the overall response to phenol in *B. subtilis* does not appear to be dictated by the relative abundance of its chemoreceptors. It is possible that a different mechanism, such as the biphasic adaptation response described for indole, is at play (57). In particular, *E. coli* cells previously adapted to roughly 700 µM indole showed an attractant response to high indole concentrations (2 mM) in chemotaxis assays while a repellent response was seen for unprimed cells. However, *B. subtilis* cells were not previously adapted to phenol in our capillary assay experiments. One possibility is that the overall response depends on the kinetics and conformational changes induced upon binding of phenol to the three chemoreceptors. Unfortunately, capillary assays are limited in quantifying the exact effect of phenol addition or removal. For example, we have previously found that the chemoeffector concentrations experienced by the cells near the capillary are 10-50 times lower than the initial concentration inside the capillary (25). The attractant and repellent capillary assays are also optimized at different cells and chemoeffector concentrations, which limits direct comparison of opposing responses induced by a chemoeffector. The mechanism of inversion of attractant to repellent response is a topic of ongoing study.

Phenolic compounds are ubiquitous in nature (58). Plant root exudates contain phenolic compounds that act as signaling molecules for plant-microbe interactions (59, 60). Chemotaxis to plant root exudates has been shown to be important for root colonization (61). *B. subtilis* is a member of plant-growth promoting rhizobacteria (62). Root exudates from soybean and rice plants are known to attract *Bacillus amyloliquefaciens* and *Bacillus spp* (16, 63). Chemotaxis was also found to be essential for early root colonization of *Arabidopsis thaliana* by *B. subtilis* (64). This study also reported an increase in the chemotaxis response to root exudates by a Δ*mcpA* strain. As McpA was found to be the major repellent chemoreceptor for phenol and other aromatic compounds in this study, the increased chemotaxis response in the Δ*mcpA* strain can be explained as due to loss of the repellent chemoreceptor. Other compounds including p-hydroxybenzoic acid, vanillyl alcohol, and isoflavones are known chemoattractants for *Agrobacterium* and *Rhizobium* species of *Rhizobiaceae* family (60). Certain phenolic compounds such as salicylic acid and hydroxycoumarins are also produced by plants in response to pathogen attacks (59). Salicylic acid was reported to be a repellent for *B. amyloliquefaciens* (16). The presence and concentration of phenolic compounds in root exudates thus changes according to many factors. Changing attractant and repellent chemotaxis responses to aromatics at different concentrations likely helps *B. subtilis* navigate altering rhizosphere conditions. Some yeasts such as *Saccharomyces cerevisiae* and *Candida albicans* also release phenolic compounds as quorum signaling molecules (65, 66). Aromatic compounds have also been detected in culture supernatants of *Pseudomonas fluorescens* (67). *B. subtilis* exhibits complex predator-prey relationships with many microorganisms and are known to release many antifungal and antibiotic compounds (68-70). Therefore, chemotaxis response to aromatics may also help *B. subtilis* find prey and evade predators.

Phenolic compounds are also classified as pollutants and cause various harmful effects in humans and animals. These compounds are primarily found in industrial wastes, forest fires, and are released to the environment through anthropogenic activities. Bioremediation strategies based on microbial decomposition of phenolic compounds are being used widely to remove phenols from wastewater (71, 72). Chemotaxis towards environmental pollutants can be exploited to guide bacteria and locate pollutants in the environment. As more reports of novel bacterial species responding to pollutants are being reported (26, 73), understanding of underlying mechanisms of chemotaxis is essential to enable design of new strategies facilitating bioavailability of pollutants for degradation.

## ACKNOWLEDGEMENT

We thank Dr. Lingyang Zhu, Dr. Dean Olson, and the SCS NMR laboratory at the University of Illinois at Urbana-Champaign for valuable inputs and help with NMR measurements. This work was partially funded by National Institutes of Health Grant GM054365 and by the University of Illinois through the Robert W. Schaefer Faculty Scholar fund.

## MATERIALS AND METHODS

### Chemicals, media, and growth conditions

*B. subtilis* strains were routinely grown on tryptose blood agar base plates (TBAB: 1% tryptone, 0.3% beef extract, 0.5% NaCl, and 1.5% agar) at 30 °C for 16 h. Chemotaxis experiments were performed with capillary assay minimal media (CAMM: 50 mM potassium phosphate buffer (pH 7.0), 1.2 mM MgCl_2_, 0.14 mM CaCl_2_, 1 mM (NH_4_)_2_SO_4_, 0.01 mM MnCl_2_, and 42 µM ferric citrate). Chemotaxis buffer consists of 10 mM potassium phosphate buffer (pH 7.0), 0.14 mM CaCl_2_, 0.3 mM (NH_4_)_2_SO_4_, 0.1 mM EDTA, 5 mM sodium lactate, and 0.05% (v/v) glycerol. All aromatic compounds were reagent grade and above. Chemicals were purchased from Sigma and Fisher.

### Strains and plasmids

All *B. subtilis* strains were derived from the chemotactic strain OI1085 (74). The strains and plasmids used in this work are listed in **Tables 2** and **3**, respectively. All cloning was performed using NEB® 5-alpha Competent *E. coli* (New England Biolabs).

**TABLE 2.**
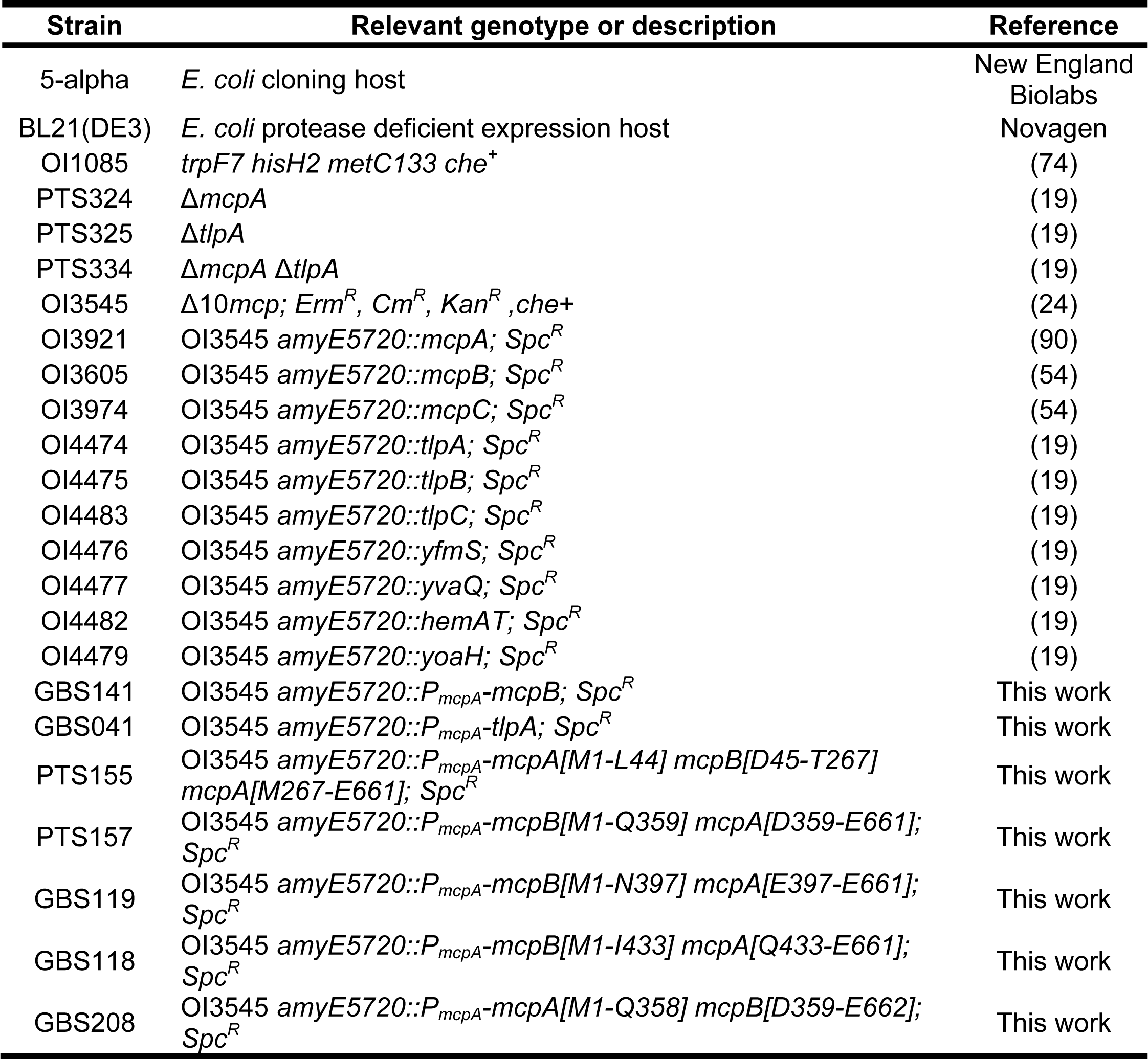
Strains used in this study.

**TABLE 3.** Plasmids used in this study.

| Plasmid | Description | Reference |
| --- | --- | --- |
| pET28a (+) | His-tagged cloning vector for protein purification; Kan<sup>R</sup> | Novagen |
| pAIN750 | *B. subtilis* empty vector for integration at *amyE*; Amp<sup>R</sup>, Spc<sup>R</sup> | (90) |
| pAIN750*mcpA* | pAIN750::*mcpA* | (54) |
| pAIN750*mcpB* | pAIN750::*mcpB* | (54) |
| pPT059 | pAIN750::*mcpA*[M1-L44] *mcpB*[D45-T267]<br>*mcpA*[M267-E661] | This work |
| pPT086 | pAIN750::*mcpA*[M1-Q358] *mcpB*[D359-E662] | (25) |
| pPT061 | pAIN750::*mcpB*[M1-Q359] *mcpA*[D359-E661] | This work |
| pGB43 | pAIN750::*mcpB*[M1-N397] *mcpA*[E397-E661] | (25) |
| pGB34 | pAIN750::*mcpB*[M1-I433] *mcpA*[Q433-E661] | (25) |
| pGB131 | 6xHis-N terminal *McpB* expression plasmid,<br>pET28(a)::*mcpB<sub>C</sub>* | This work |
| pGB130 | 6xHis-N terminal *McpA* expression plasmid,<br>pET28(a)::*mcpA<sub>C</sub>* | This work |
| pPT021 | 6xHis-C terminal *McpA* expression plasmid,<br>pET28(a)::*mcpA<sub>S</sub>* | This work |
| pGB46 | 6xHis-N terminal *HemAT* expression plasmid,<br>pET28(a)::*hemAT<sub>C</sub>* | (25) |
| pSP03 | 6xHis-N terminal *HemAT* expression plasmid,<br>pET28(a)::*hemAT<sub>S</sub>* | (25) |

Receptor chimeras were constructed according to the procedure described previously (25). Briefly, the region outside the fusion points was amplified by whole plasmid PCR with the pAIN750*mcpA* plasmid as the template. The desired fragments from *mcpB* were amplified from pAIN750*mcpB* with a short overlap on both ends. The PCR products were purified after gel extraction, assembled using Gibson assembly (75), and transformed in *E. coli*. After sequence verification, the correct plasmids were linearized at the *Xho*I restriction site, re-ligated, and transformed into *B. subtilis* OI3545 (receptorless mutant) using the two-step Spizizen method (76). Colonies were screened for spectinomycin resistance and integration in the *amyE* locus was verified using Gram iodine solution (0.33% iodine, 0.66% potassium iodide, and 1% sodium bicarbonate) on starch plates. Correct clones were unable to hydrolyze starch and form clear zones.

For construction of chimeric receptors under the control of the *mcpA* promoter, whole plasmid PCR was used to amplify the pAIN750*mcpA* plasmid excluding the *mcpA* region to be swapped by the desired region from another receptor. The desired region from another receptor was also PCR-amplified with overlapping primers and both DNA fragments were purified and assembled with Gibson assembly. Point mutations were performed as described previously (25).

Cloning for recombinant protein production was performed in *E. coli* BL21(DE3). The DNA fragments corresponding to the signaling domain of McpA (residues 359 to 661) and McpB (residues 359 to 662) were PCR-amplified and cloned in frame with a N-terminal His_6_-tag in pET28a(+) plasmid at *NheI* restriction site using Gibson assembly. McpA sensing domain (residues 21 to 278) was cloned in frame with a C-terminal His_6_-tag in pET28a(+) plasmid between the *Xho*I and *Nco*I restriction sites. McpC sensing domain (residues 33 to 278) was cloned in frame with a N-terminal His_6_-tag in pET28a(+) at the *Nde*I restriction site using Gibson assembly. Assembled plasmids were transformed in *E. coli*.. After isolation and sequence verification, all plasmids were transformed into *E. coli* BL21(DE3) strain for protein expression and purification.

### Capillary assay for chemotaxis

The “chemical-in-pond” modification of the capillary assay was used for quantifying repellent chemotaxis responses (PMID: 4597449). Briefly, *B. subtilis* strains were grown for 16 h at 30 °C on TBAB plates. The cells were scraped from the plates and resuspended to *OD*_600_ = 0.03 in 5-mL CAMM supplemented with 50 μg/mL histidine, 50 μg/mL methionine, 50 μg/mL tryptophan, 20 mM sorbitol, and 2% TB. The cultures were then grown to *OD*_600_ = 0.4 – 0.45 at 37 °C with shaking at 250 rpm, after which 50 μL of GL solution (5% (v/v) glycerol and 0.5 M sodium lactate) was added, and cells were incubated for another 15 min. The cells were then washed twice with chemotaxis buffer and incubated for additional 25 min at 37 °C with shaking at 250 rpm to assure that the cells were motile. Cells were then diluted to *OD*_600_ = 0.01 in chemotaxis buffer containing appropriate concentrations of repellents. The culture was shaken for 10 min at room temperature, shaking at 150 rpm and then aliquoted into 0.3-mL ponds on a slide warmer at 37 °C. Closed-end capillary tubes filled with chemotaxis buffer were inserted in the ponds. After 1 h, cells in the capillaries were harvested and transferred to 3 mL of top agar (1% tryptone, 0.8% NaCl, 0.8% agar, and 0.5 mM EDTA) and plated onto TB agar (TB and 1.5% agar) plates. These plates were incubated for 16 h at 37 °C and colonies were counted. Experiments were performed in triplicate each day and repeated on three different days. Attractant assays for aromatic compounds were carried out at *OD*_600_ = 0.001 for 1 h. For testing functionality of the mutant strains expressing chimeric receptors, attractant assays were performed for 30 min with cells diluted to *OD*_600_ = 0.001 in the pond and asparagine solution (3.16 μM) in the capillary. For measuring chemotaxis response to external acidic pH, capillaries filled with chemotaxis buffer at pH 7.0 were inserted in the pond containing cells at *OD*_600_ = 0.001 preadapted to pH 8.0 in chemotaxis buffer and cells in the capillaries were harvested after 1 h and counted as described above (19).

### Protein purification

*E. coli* BL21(DE3) cells harboring the His_6_-tagged expression plasmids were grown in 2 L flask containing LB medium supplemented with 30 µg/mL kanamycin at 37 °C and shaking at 250 rpm until *A*_600_ = 0.7 at which point expression was induced by the addition of 1 mM Isopropyl β-D-1-thiogalactopyranoside (IPTG). The cultures were grown for 12 h at 25 °C and then harvested by centrifugation at 7,000 x *g* at 4°C for 10 min. Cells were resuspended in lysis buffer (50 mM NaH_2_PO_4_, 300 mM NaCl, 10 mM Imidazole, pH 8) and sonicated (5 x 10 s pulses). After centrifugation at 40,000 x *g* for 1 h, the supernatant was loaded on a 5 mL GE Hi-Trap Chelating column charged with 0.1 M NiSO_4_ and binding buffer (50 mM NaH_2_PO_4_, 300 mM NaCl, 20 mM imidazole, pH 8). The protein-bound column was then washed with 10 column volumes of binding buffer and proteins were eluted with imidazole gradient of 20-500 mM. The collected protein fractions were pooled and concentrated using an Amicon ultrafiltration cell (Millipore) and dialyzed into PBS (10 mM Na_2_HPO_4_, 1.8 mM KH_2_PO_4_, 137 mM NaCl, 2.7 mM KCl, pH 7.4) at 4 °C. HemAT domains were purified as described before (25). Aliquots were stored at -80°C. Concentration of purified proteins was measured by the Pierce BCA protein assay kit.

### Saturation-transfer difference nuclear magnetic resonance spectroscopy (STD-NMR)

All NMR spectroscopy measurements were performed on a Varian VNMRS instrument at 750 MHz with 5 mm Varian HCN probe at 298 K without sample spinning. Prior to measurements, protein samples were buffer exchanged into PBS (50 mM KH_2_PO_4_, 20 mM NaCl, pH 7.4) in D_2_O using Micro Bio-Spin® Columns with Bio-Gel® P-6 (Bio-Rad Laboratories). Protein (final concentration 20 µM) and ligand samples (final concentration 2 mM) were added to the NMR tube at 500 µL solution volume. For K_d_ measurements, ligand concentrations were varied from 0.5 mM to 4 mM. ^1^H spectra were obtained from 32 scans with a 90-degree pulse and a 2-s relaxation delay. In STD-NMR experiments, the protein samples were selectively saturated at 0.5 ppm with a train of Gaussian pulses of 50 ms duration with 0.1 ms delay and 5 s relaxation delay for a total saturation time of 1-4 s and 256 scans. Off-resonance irradiation was applied at 30 ppm. Trim pulse of 50 ms was used to reduce protein background. All STD spectra were obtained by phase cycling after a block size of 8 to reduce artifacts resulted from temperature variation and magnet instability. Manual subtraction was performed to obtain the difference spectra. Control experiments were performed on samples containing only ligands without protein. All areas were calculated using MNova V14.1 (Mestrelab chemistry solutions) in stacked mode.

The STD amplification factor (STD-AF) is defined as the fractional saturation of a given proton multiplied by the excess of the ligand over the protein (38).

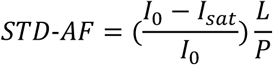

The method of initial slopes was used to determine the dissociation constant (K_d_) (38, 77). The STD amplification factors at initial slopes (STD-AF_0_) for each ligand concentration were calculated by plotting the STD-AF evolution with saturation times and fitting the equation:

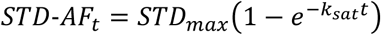

The initial slope was then obtained by taking the derivative at t=0, yielding:

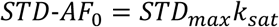

Plotting STD-AF_0_ values against the increasing ligand concentrations (L) would yield a Langmuir hyperbolic dose-response curve. Finally, the dissociation constant (K_d_) was calculated by fitting the data to the following equation:

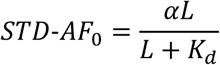

### Phylogenetic tree and structural analyses of chemoreceptor domains

*B. subtilis* (RefSeq: GCF_000009045.1) chemoreceptor sequences were obtained from MiST 3.0 database (78). Tree analysis was carried out using TREND platform (79). Protein sequences were first aligned with the online version of MAFFT (80) and the alignment result was then used as an input in FastTree program (81) to construct the tree. Domain predictions was performed using HMMER3 (82).

### *In-silico* docking experiments

The dimer structure of McpA signaling domain (residues 352 to 661) was constructed using Modeller (v-9.23) (83). The homology model was based on *Thermotoga maritima* Tm113 chemoreceptor (PDB 2CH7) (84). Side-chain conformations were refined using SCWRL4 (85) and the resulting protein structure was minimized using YASARA server (86). 3D structures of the ligands were obtained from Pubchem (87). Docking experiments were carried out using Autodock Vina (v-1.1.2) (88) in which the grid size was set to 58 x 50 x 62 points with 1 Å spacing surrounding residues 391 to 436 and exhaustiveness was set at 10. Clusters were visualized using Autodock Tools GUI (89).

## Data availability

Raw data are provided as Supplemental Dataset 1.

## ACKNOWLEDGEMENTS

This work was supported by the University of Illinois through the Robert W. Schaefer Faculty Scholar fund.

